# Structural insight into the putative role of novel SARS CoV-2 E protein in viral infection: a potential target for LAV development and therapeutic strategies

**DOI:** 10.1101/2020.05.11.088781

**Authors:** Manish Sarkar, Soham Saha

**Affiliations:** Structural Biology Lab, Department of Biochemistry, Bose Institute, P-1/12, CIT Road Scheme VIIM Kolkata 700054 WB India; Laboratory for *Perception and Memory*, Institut Pasteur, F-75015 Paris, France; Centre National de la Recherche Scientifique (CNRS), Unité Mixte de Recherche (UMR-3571), F-75015 Paris, France

**Keywords:** SARS-CoV-2, COVID-19, E-protein, homology modeling, proton channel

## Abstract

The outbreak of COVID-19 across the world has posed unprecedented and global challenges on multiple fronts. Most of the vaccine and drug development has focused on the spike proteins and viral RNA-polymerases. Using the bioinformatics and structural modeling approach, we modeled the structure of the envelope (E)-protein of novel SARS-CoV-2. The E-protein of this virus shares sequence similarity with that of SARS-CoV-1, and is highly conserved in the N-terminal regions. Incidentally, compared to spike proteins, E proteins demonstrate lower disparity and mutability among the isolated sequences. Using homology modeling, we found that the most favorable structure could function as a gated proton channel. Combining pocket estimation and docking with water, we determined that GLU 8 and ASN 15 in the N-terminal region were in close proximity to form H-bonds. Additionally, two distinct “core” structures were visible, the hydrophobic core and the central core, which may regulate the opening/closing of the channel. We propose this as a mechanism of viral proton channeling activity which may play a critical role in viral infection. In addition, it provides a structural basis and additional avenues for LAV development and generating therapeutic interventions against the virus.

**Significance Statement:** Structural modeling of the novel SARS-CoV-2 envelope protein (E-protein) demonstrating its possible proton channeling activity

## Introduction

The COVID-19 (CoronaVIrus Disease 2019) is a severe acute respiratory syndrome (SARS) caused by a novel coronavirus, SARS-CoV-2, and has taken the form of a worldwide pandemic in the last few months (1). To date, this disease has affected nearly 4 million people, resulting in nearly 300 thousand deaths disrupting social and economic structures of nearly 190 nations across the globe with numbers still on the rise (2, 3).

Coronavirus is a positive-sense single-stranded RNA virus belonging to the class of β-coronaviruses of the family *Coronaviridae*, and affected humans causing different forms of mild common cold. Coronaviruses like hCoV-OC43, HKU, and 229E have been responsible for these diseases (4). The emergence of human coronaviruses in the 21st century with enhanced pathogenicity was seen in the case of SARS-CoV-1 mediated infection in 2002, reporting almost 8000 detected cases worldwide and a mortality rate of 10 %. This was followed by MERS-CoV mediated near-pandemic in 2012 infecting almost 2500 people worldwide with a mortality rate of 36% (5). The ongoing novel SARS-CoV-2 pandemic, on the other hand, has a lower fatality rate in comparison with the previous coronavirus outbreaks, but a higher human-to-human contagion efficiency than the previous ones (6). This has led to the spread of this infection worldwide, across continents, affecting millions.

Coronaviruses have four main structural proteins- (i) Nucleocapsid protein (N), (ii) Spike protein (S), (iii) Membrane protein (M), and (iv) Envelope protein (E). The E protein is the smallest of all the structural proteins of 8-12 kDa and is involved in a wide spectrum of functional repertoire (7). It comprises of three domains: (i) short hydrophilic N-terminus domain consisting of 7-12 amino acids, (ii) hydrophobic transmembrane domain which is around 25 amino acids long, and (iii) long hydrophilic C terminal (8–10). The hydrophobic transmembrane domain oligomerizes to form pentameric ion channels in the ERGIC (ER-Golgi Intermediate compartment) membrane having low or no ion selectivity (11–13). A β-coil-β motif present in the C terminal of E protein contains a conserved proline residue (14) which is indispensable for its localization in the ER-Golgi complex (15). Previous studies show that the last 4-5 residues of SARS-CoV E protein form a PDZ-binding motif, deletion of which resulted in revertant mutants and reversal of its pathogenicity (16, 17). This, interestingly, points to a potential target for the development of live attenuated vaccines (LAVs): generating certain mutations in the E protein while keeping the PDZ-binding motif intact (7). During the viral replication cycle, the virus incorporates a small proportion of the viral E protein expressed within the infected host, into the virion particles (18). The larger proportion of E protein is localized at the sites of mammalian intracellular trafficking, the ER-Golgi network and the ERGIC (19). This localization in the intracellular trafficking components of the host cell helps in the assembly of viral structure and budding (19). Coronaviruses lacking the E protein produced virion particles with crippled maturation and were incompetent in forming new virion particles (20–24). The role of the E protein as a non-specific ion channel on the ER-Golgi membrane of the host cell has also been suggested (12, 13, 19). Recent studies have shed light on a large number of host-pathogen protein-protein interactions (PPIs) and identified two classes of small molecules - mRNA translation inhibitors and Sigma factor 1 and 2 receptor regulators, which showed antiviral activity (5) against novel SARS-CoV-2.

While there are no antiviral drugs or effective vaccines against COVID-19 on the market; we leverage the existing knowledge of viral channel proteins and structural modeling to explain a putative mechanism involving E protein in SARS-CoV-2 related infection. The development of therapeutic approaches is being hindered due to the lack of understanding of molecular mechanisms and molecular players responsible for this novel viral infection (5). The possibility of E protein being a vaccine candidate has already been explored with the MERS-CoV; certain mutations towards the C-terminal end combined with mutations in the NS1 protein resulted in a stable vaccine against the virus, overcoming the pathogenesis of the revertant mutants (16). Since CoV E-protein forms pentameric non-selective ion channels in the membrane (11–13), we modeled the SARS-CoV-2 E protein using the SARS-CoV-1 NMR structure (10) as the template. M2 viroporin of Influenza A virus has been reported to function as a proton channel with a water-hopping mechanism in a conformational dependent manner (25). Similarly, we identified key amino acid residues lining the inner luminal side of the SARS-CoV-2 E protein, and subsequent water docking models suggest that the pentameric structure is able to form dynamic closed and open states. While it calls for detailed studies, the E-protein presents a potential target for the development of live attenuated vaccines (LAVs) and other therapeutic strategies to curb the COVID-19 pandemic.

## Results

### Coronavirus E-proteins are conserved and have lower mutability compared to spike proteins

A BLASTp search run resulted in 54 sequences of E-proteins from different origins (**Fig. 1a**). E protein of SARS-CoV-2 isolated in April (Accession number: QII57162.1) was used as the search reference. Some E-proteins specific to bat (HKU3-7), human SARS 2018, bat SARS_RsSHC014, bat BtRI-BetaCoV, human SARS-CoV-1 and human SARS-CoV-2 (Accession number: QII57162.1) show a high degree of conservation in their amino acid composition (**Fig. 1b**). In comparison, we also looked at the conservation of amino acids for the spike proteins (**Suppl. Fig. 1a**), M-proteins (**Suppl. Fig. 1b**) and protein 7a (**Suppl. Fig. 1c**). The candidate E-proteins from different origins were closely associated when we plotted the evolutionary distance (**Suppl. Fig. 1d**). All the selected sequences of Spike proteins, M proteins, E-proteins, and protein 7a were selected based on a 90-100% sequence identity. The amino acid composition indicated that the E-proteins have a larger proportion of Leucine (LEU) and Valine (VAL) compared to either the Spike or the M-proteins (**Fig. 1c i-iii**). Of particular interest was the increased disparity index among spike proteins compared to the E-proteins (**Fig. 1d i-ii)**. The value of the disparity index close to 1 (red) indicated larger differences in amino acid composition bias than expected only out of chance probability and evolutionary divergence (26). Using the amino acid composition, we calculated the probability of accumulating mutation for a particular residue as a proxy of mutability. We found that leucine (LEU, 0.189 ± 0.132), valine (VAL, 0.25 ± 0.13) and serine (SER, 0.461 ± 0.091) (**Fig. 1e;** indicated by $$) of the E proteins have lower than 50% chance of mutability compared to spike proteins (LEU (0.626 ± 0.004), VAL (0.652 ± 0.004) and SER (0.632 ± 0.006)) (**Fig. 1e**). Given the role of E proteins in viral infection (19), we think that E-proteins present to us a relatively stable viral protein that demands greater investigation to elucidate its mechanistic insights and features.

**Figure 1.**
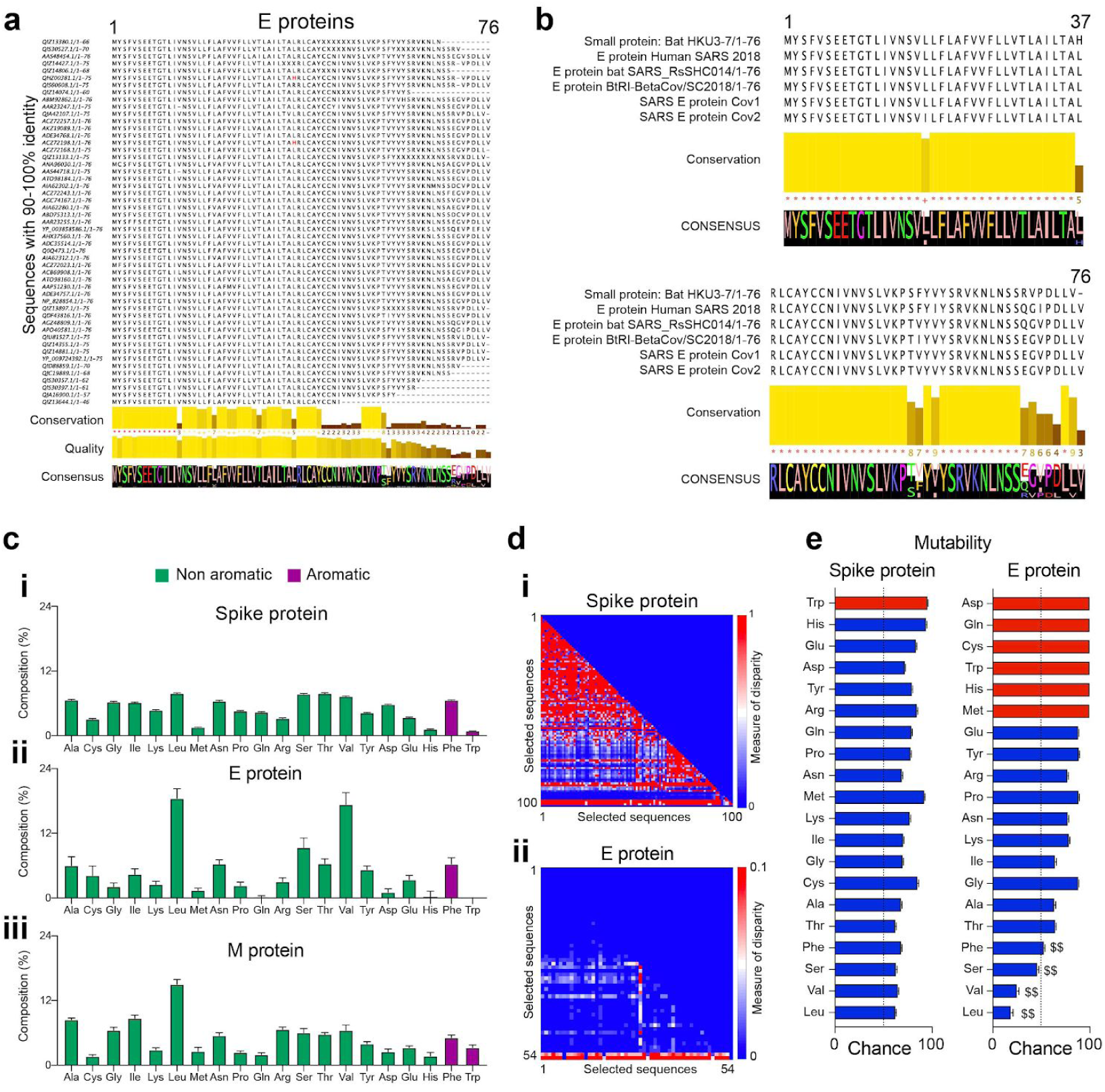
**(a)** Sequence alignment of CoV E-proteins performed by Clustal Omega after a BLASTp search against human SARS-CoV-2 E protein (Accession number: QII57162.1). The selected aligned sequences were based on the criteria of 90-100% similarity. The conserved sequences, quality of sequence alignment, and consensus sequence are shown below. **(b)** Sequence alignment of CoV E-proteins from the bat (HKU3-7), human SARS 2018, bat SARS_RsSHC014, BtRI-BetaCoV, SARS-CoV-1 and SARS-CoV-2 showing highly conserved regions in the N-terminal region. The consensus sequence is shown below. **(c)** Percentage composition of each amino acid computed from the aligned selected sequences for (i) Spike protein, (ii) E-protein, and (iii) M-protein. **(d)** Lower triangular heatmap representation of the disparity index computed from the sequence alignment for (i) spike protein, and (ii) E-protein. Scale: 0-1 (for i), and 0-0.1 (for ii). **(e)** Mutability (probability of amino acid change) calculated for spike protein and E-protein. The dotted line indicates a 50% probability chance, while red bars indicate amino acids absent/single-site presence on the sequences analyzed. $$ indicates residues with lower mutability in E-protein compared to spike protein.

### Homology modeling of SARS-CoV-2 E protein, validation and bottleneck measurements

We designed a template-based pentameric structural model of the E protein of SARS-CoV-2 with the NMR structure of E protein from SARS-CoV-1 (PDB ID: 5×29; (10)) as the template (**Fig. 2a**). The putative pentameric models of the E-protein have been obtained from GalaxyWEB server (**Fig. 2b-i, side view, Fig. 2d-i, top view**) and SWISS-MODEL (**Fig. 2b-ii-side view, Fig. 2d-ii-top view**). Based on homology modeling, we obtained 16 different structural models from both the GalaxyWEB (**Suppl. Fig. 2a-p**) and SWISS-MODEL (**Suppl. Fig. 3a-p**). The model obtained from GalaxyWEB spanned the entire protein (1-75) but the five residues (PDLLV) in the C-terminal end of the protein were removed which reduces the structural flexibility of that region improving the MolProbity parameters. The model obtained from SWISS-MODEL spanned from 8-65 residues similar to the NMR structure. All the models from both these protocols and the template structures were validated using MolProbity. The GalaxyWEB models were chosen based on the following observations: (i) they spanned a longer region of the protein **(Fig. 2g-i, ii)** and (ii) they had better validation scores (Clash score and MolProbity score). The final structural model (**Fig. 2b-i)** was chosen depending on the structural validation parameters obtained from MolProbity **(Fig. 2h-green**). We refer to the structural model used here for further characterization as “*E-put*”. The bottleneck radius of the template structure (4.2Å; **Fig. 2e** and the models obtained from GalaxyWEB (3.4Å; **Fig. 2f-i)** and SWISS-MODEL (3.4Å; **Fig. 2f-ii)** were calculated using the centroid formed by the triangular orientation of the PHE 26 residues **(Fig. 2h;** refer to *Methods*).

**Figure 2.**
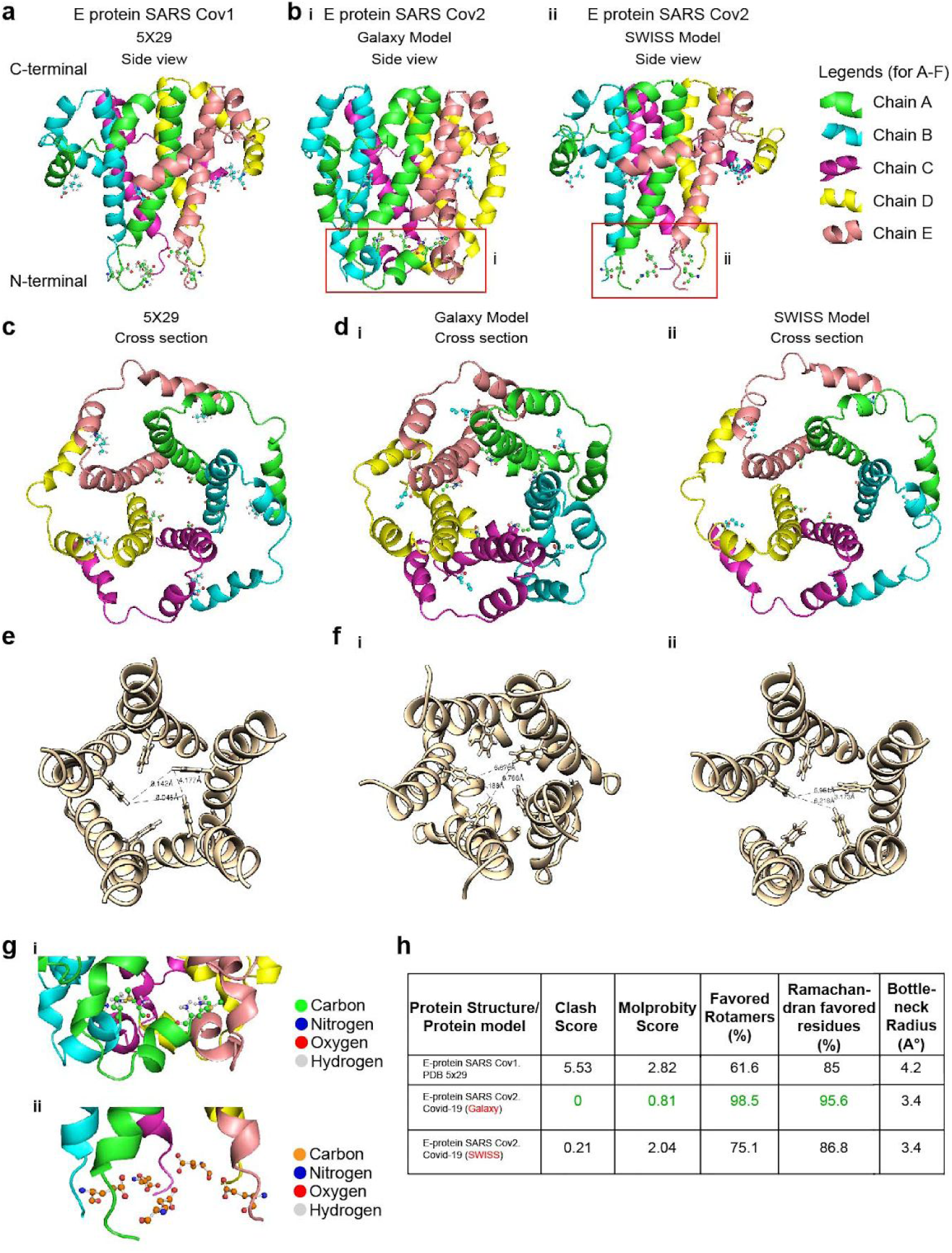
**(a)** NMR structure of pentameric E protein (PDB id: 5×29) of SARS-CoV-1. **(b)** Pentameric model of E protein of SARS-CoV-2 (COVID-19) (i) generated in GALAXY (*E-put*), and (ii) generated in SWISS-MODEL. A magnified figure for each is provided in (red box). **(c)** Top view of pentameric E protein SARS-CoV-1 (5×29). **(d)** Top view of pentameric E protein SARS-CoV-2 (i) generated in GALAXY (*E-put*), and (ii) generated in SWISS-MODEL. **(e)** Bottleneck radius measurement of C (5×29). **(f)** Bottleneck radii measurement of D (i) GALAXY (*E-put)* and (ii) SWISS-MODEL. **(g)** N-terminal residues of (i) GALAXY model (*E-put)* (as in B-i) and (ii) SWISS-MODEL model (as in B-ii). **(h)** Structure validation parameters and pore radii of 5×29, GALAXY model (*E-put)*, and SWISS-MODEL model.

### *In silico* structural analysis of *E-put* showing water docking and putative lining residues important for its functionality as a proton channel forming H-bonds with water

The pore volume of the *E-put* protein was calculated using the CASTp server clearly showing a discontinuous channel in the luminal surface of the pore consisting of Pocket 1 and Pocket 2 **(Fig. 3a).** The lining residues were LEU 37, ILE 33, VAL 29, PHE 26, ALA 22, LEU 19, LEU 18, ASN 15, GLU 8, VAL 5 and PHE 4 **(Fig. 3b-i).** The luminal surface of the pore is shown according to its hydrophobicity content **(Fig. 3b-ii).** The presence of pockets lined by charged, aromatic and hydrophobic residues suggested that the pentamer could possibly form continuous water chains for proton channeling (25). To gain an insight into this hypothesis, we docked *in silico* water molecules with the *E-put* protein using SWISS-DOCK. The presence of water molecules docked in the N-terminal end was observed **(Fig. 3c).** The distances of the marked residues (LEU 37, ALA 36, ILE 33, ASN 15 and GLU 8) from their nearest water molecules showed that ASN 15 and GLU 8 lie in the range of hydrogen bonding limit (<2.5 Å; **Fig. 3d).** As expected, the distance of LEU 37, ALA 36, ILE 33 from the nearest water molecules lie beyond the hydrogen bond limit and also short-range interactions limit (>3.8 Å; **Fig. 3d).**

**Figure 3.**
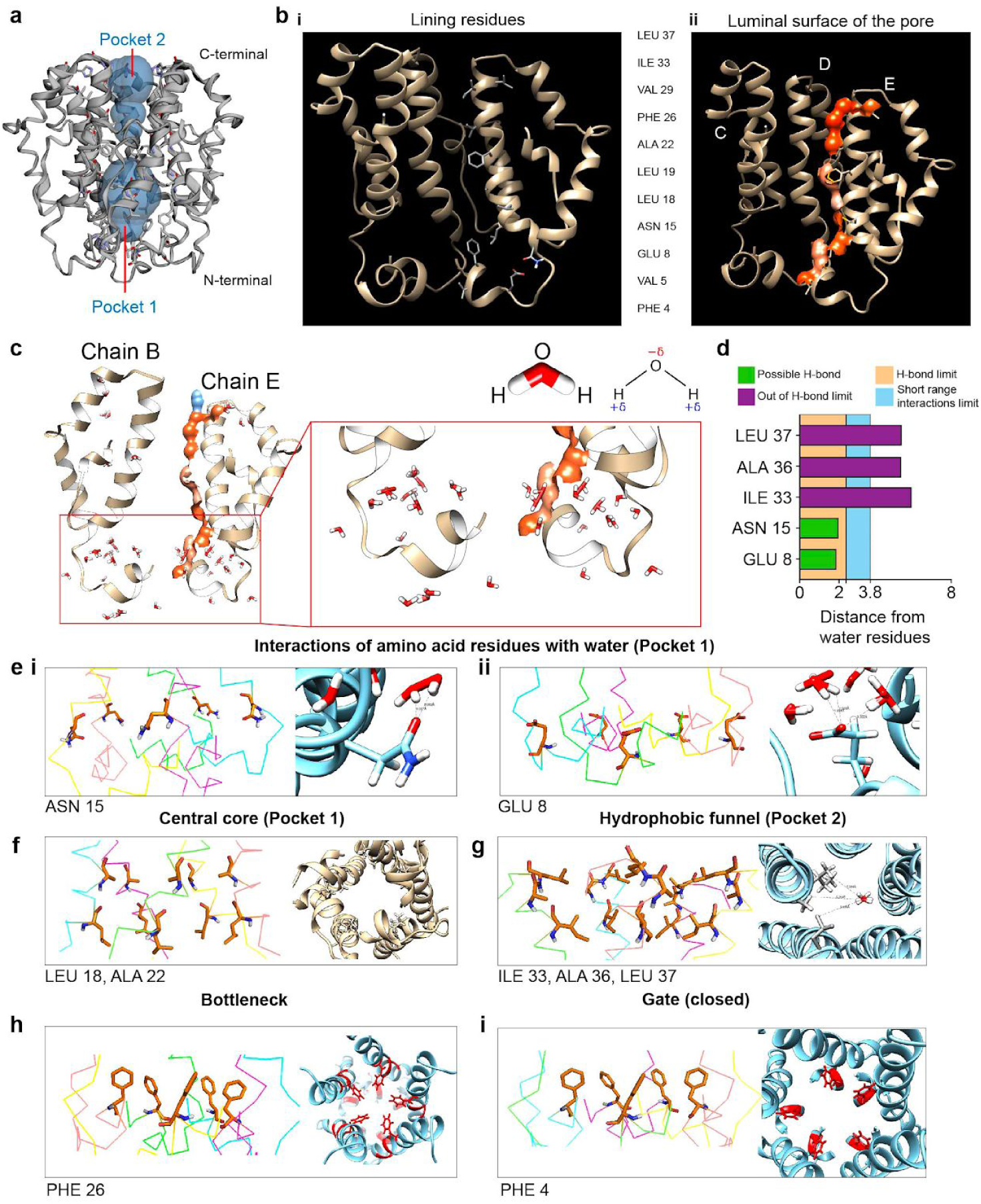
**(a)** Estimation of the pore volume of the *E-put* protein using the CASTp 3.0 server. **(b)** (i) Residues lining the luminal side of the *E-put* protein core and (ii) the luminal surface of the pore determined by the CASTp 3.0 server. **(c)** Docking of water to the *E-put* protein by SWISS-DOCK showing the chains B and E of the pentamer and magnification of the region of interaction. **(d)** Distances of water molecules from different lining residues and the limit of H-bond. **(e)** Residues’ orientation of the *E-put* protein generated from PYMOL and interaction of water with residues showing the distance between water and (i) ASN 15 and (ii) GLU 8 by CHIMERA. **(f)** Residues’ orientation generated from PYMOL (side view) and CHIMERA (top view) lining the Central core of the *E-put* protein. **(g)** Residues’ orientation generated from PYMOL and interaction of water with the lining residues of the Hydrophobic funnel of the *E-put* protein by CHIMERA. **(h)** Residues’ orientation generated from PYMOL (side view) and CHIMERA (top view) showing the Gate of the *E-put* protein in the closed conformation. **(i)** Residues’ orientation generated from PYMOL (side view) and CHIMERA (top view) forming the bottleneck of the *E-put* protein.

The structural orientation of the ASN 15 and GLU 8 of the *E-put* protein made these residues ideal candidates for water docking **(Fig. 3e-i-ii, left, side view, Suppl. Fig. 4a-i-ii, top view).** The distance from a dimer water molecule docked in the luminal side of the pore to the carbonyl group of the amide side chain of ASN 15 and the charged carboxyl group of the side chain of GLU 8 was 2.03Å and 1.94Å respectively **(Fig. 3e-i-ii, right).** Both were seen to be perfectly within conventional H-bonding limits.

The structural orientation of LEU 18 and ALA 22 along with their side chains formed the ‘central core’ (Pocket 1) of the *E-put* protein **(Fig. 3f, left, side view, Fig. 3f, right, top view, Suppl. Fig. 4b, top view).** The LEU 37, ALA 36 and ILE 33 residues along with their side chains formed the ‘hydrophobic funnel’ (Pocket 2) of the *E-put* protein **(Fig. 3g, left, side view, Suppl. Fig. 4c, top view).** The distance from a trimer of water molecules docked in the luminal side of the pore to the hydrophobic side chains of the comprising residues was much higher than H-bonding limit (2.5 Å) and short-range interactions limit (3.8 Å) **(Fig. 3g, right;** LEU 37-5.39 Å, ALA 36-5.33 Å, ILE 33-5.91 Å). This shows that water molecules can access the interior of the hydrophobic funnel without any unfavorable interactions or steric clashes.

The structural orientation of PHE 26 and their side chains formed the bottleneck of the *E-put* protein, which can potentially limit the conduction of water or ions through the channel (**Fig. 3h, left, side view, Fig 3d, right, top view**). The structural orientation of PHE 4 forming the gate of the *E-put* protein, the conformation of which could determine the putative closed (**Fig. 3i, left, side view, Fig 3i, right, top view**) and open states (**Suppl. Fig. 4d, left, side view, Fig Suppl. Fig. 4d, right, top view**).

Simultaneously, we analyzed the model obtained from SWISS-MODEL **(Fig 2-b-ii, d-ii).** CASTp pocket analysis clearly showed the discontinuous channel in the luminal surface of the pore consisting of Pocket 1 and Pocket 2 **(Fig. 4a)**, and the lining residues **(Fig. 4b-i).** The hydrophobicity content is similarly visible on the luminal surface **(Fig. 4b-ii).** We docked *in silico* water molecules with the SWISS-MODEL protein using SWISS-DOCK (**Fig. 4c)**. Similar to E-put shown above, the distances of ASN 15, and GLU 8 from their nearest water molecules showed that they lie in the range of hydrogen bonding limit (<2.5 Å; **Fig. 4d-i, ii)**, while LEU 37, ALA 36, ILE 33 lie beyond the hydrogen bond limit and also short-range interactions limit (>3.8 Å; **Fig. 4e**). The distance from a dimer water molecule to the carbonyl group of ASN 15 and the charged carboxyl group of GLU 8 was 1.95 Å and 1.88 Å respectively **(Fig. 4f**). The PHE 26 and their side chains formed the bottleneck of the SWISS-MODEL protein (**Fig. 4-g**).

**Figure 4.**
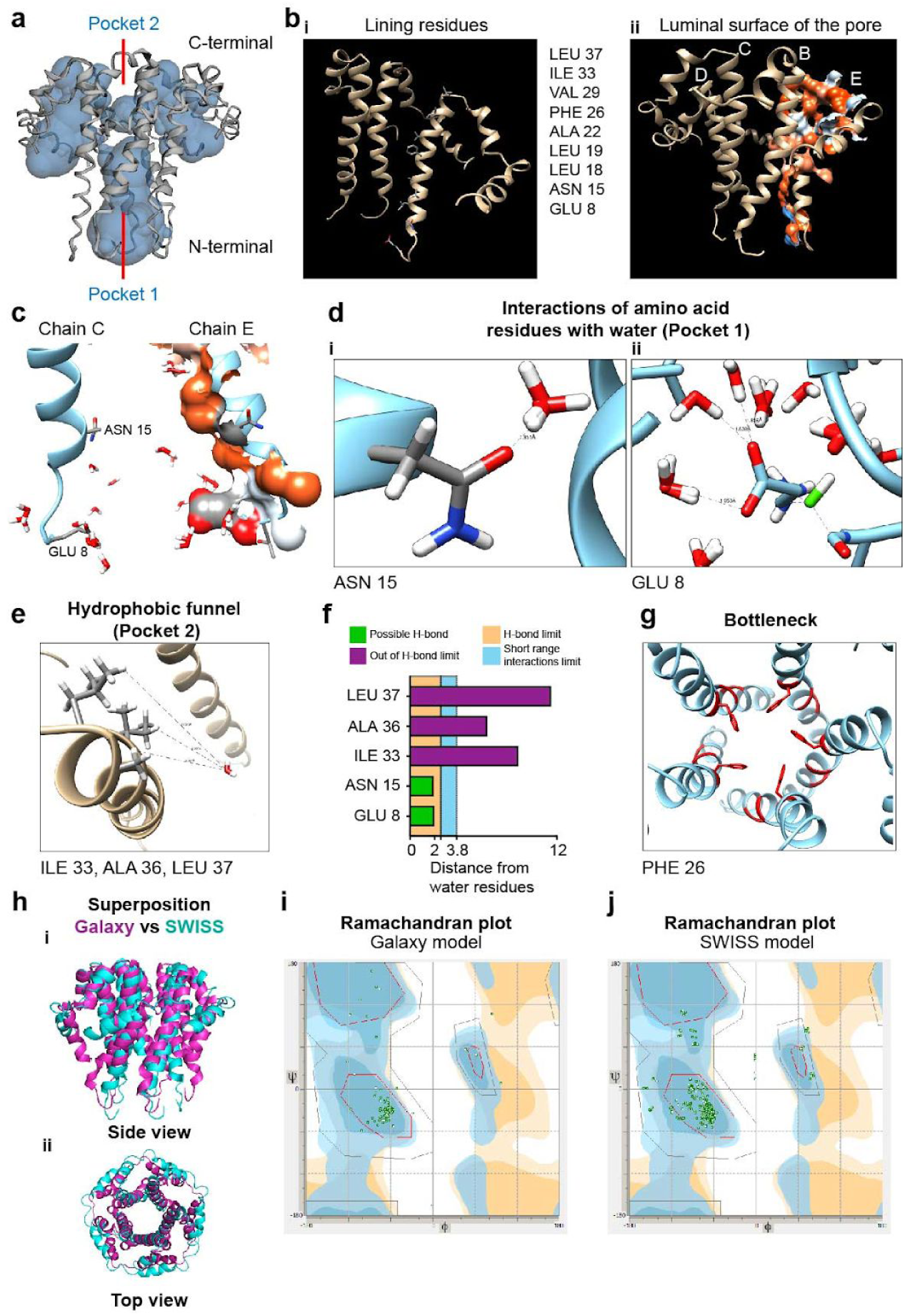
**(a)** Estimation of the pore volume of the SWISS-MODEL protein using the CASTp 3.0. **(b)** (i) Residues lining the luminal side of the SWISS-MODEL protein core and (ii) the luminal surface of the pore determined by the CASTp 3.0 server. **(c)** Docking of water to the SWISS-MODEL protein by SWISS-DOCK showing the chains C and E of the pentamer. **(d)** Residues’ interaction of water with residues showing the distance between water and (i) ASN 15 and (ii) GLU 8 by CHIMERA. **(e)** Residues’ interaction of water with the lining residues of the hydrophobic funnel of the SWISS-MODEL protein by CHIMERA. **(f)** Distances of water molecules from different lining residues and the limit of H-bond. **(g)** Residues’ orientation in CHIMERA forming the bottleneck of the SWISS-MODEL protein. **(h)** Superposition of the *E-put* and the SWISS-MODEL protein (purple: *E-put*; cyan: SWISS-MODEL) as (i) side view and (ii) top view. **(i)** Ramachandran plot of the *E-put* protein (Galaxy model). **(j)** Ramachandran plot of the SWISS-MODEL protein.

The modeled *E-put* protein from GalaxyWeb (Fig 2-b-i) and SWISS-MODEL (2-b-ii) was superimposed using Pymol which yielded an RMS (RMSF) of 3.865 (**Fig. 4-h-i, side view, Fig. 4-h-ii, top view).** In addition, we superimposed the *E-put* obtained from the GalaxyWeb and SWISS-MODEL with the reported NMR CoV-1 E protein structure (**Suppl. Fig. 5a, b: i (top view), ii (side view)**. Ramachandran plot of the *E-put* obtained from the GalaxyWeb and SWISS-MODEL showed that most of the favored residues of both the models fall in the alpha-helical region (**Fig. 4-i,j**).

### Docking of *E-put* with lipid molecules: Ceramide and phosphatidylcholine

Ceramide and phosphatidylcholine contribute to the maximum percentage of lipid composition in the ER-Golgi network and ERGIC complex of the mammalian cells including humans. Ceramide (**Fig. 5a**) and phosphatidylcholine (**Fig. 5d**) were docked to *E-put* protein using CHIMERA and Autodock Vina (**Fig. 5b, e** respectively; refer to *Methods*). The RMSD values for both the cases were zero, validating the quality check of the lipid-bound structure of *E-put*. The distances of the terminal guanidino group of the side chain of ARG 38 with the phosphoryl group of the charged headgroup of ceramide (2.32 Å; **Fig. 5c**) and the carbonyl group of the charged headgroup of phosphatidylcholine (2.29 Å, **Fig. 5f)** were calculated. The distances fall within the covalent or strong hydrogen bond limit (**Fig. 5g**). This concludes that the *E-put* protein docks to the lipid components of the ER-Golgi network and ERGIC complex effectively either by covalent or strong hydrogen bond formation.

**Figure 5.**
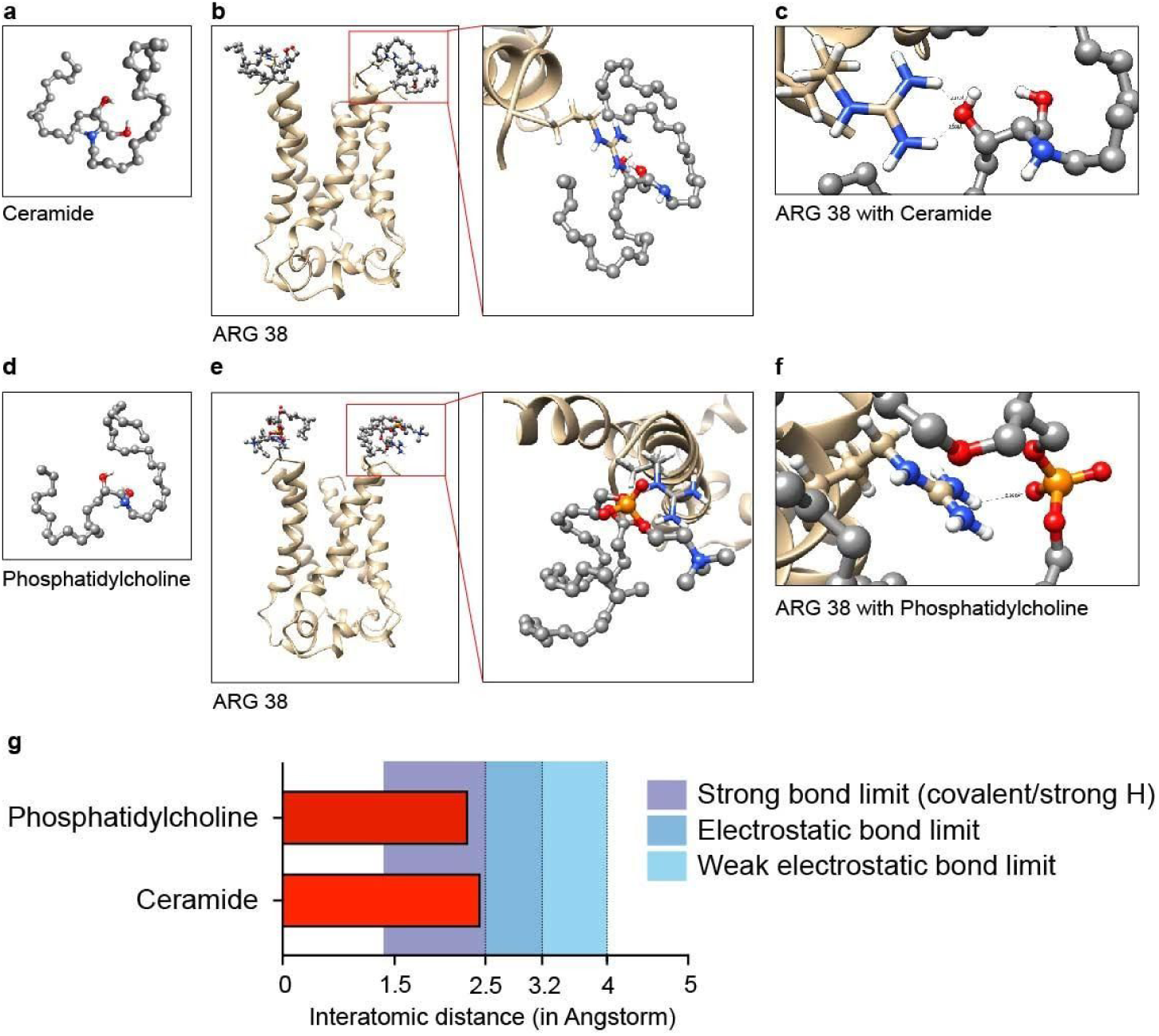
**(a)** Ceramide: Sphingolipid generated in OpenBabel and CHIMERA. **(b)** Docking of Ceramide to the N-terminus of the *E-put* protein in CHIMERA-VINA and magnification of the region of interaction. **(c)** Interaction of ARG 38 in the C terminal with the docked ligand showing the distances between the charged groups of ARG 38 and the ligand. **(d)** Phosphatidylcholine: Glycerophospholipid generated in OpenBabel and CHIMERA. **(e)** Docking of Phosphatidylcholine to the N-terminus of the *E-put* protein in CHIMERA and Autodock Vina and magnification of the region of interaction. **(f)** Interaction of ARG 38 in the C terminal of *E-put* with the docked ligand showing the distances between the charged groups of ARG 38 and the ligand in CHIMERA. **(g)** Distances of the ligands from ARG 38 showing the bond limits of different interactions.

### *In silico* structural analysis of *E-put* shows a putative conformation-dependent proton channeling mechanism

The mechanism of proton transport through the M2 channel of influenza A virus *via* the formation of well-ordered clusters of water molecules has already been reported (25). This protein undergoes a pH-dependent conformational change between its closed and open state ((25); **Fig. 6a**). Based on the M2 channel mechanism and our findings, we propose that *E-put* protein undergoes conformational changes: (1) Closed discontinuous state (**Fig. 2b-i, 6b-i, top-lateral view, bottom-cross-section**) to (2) Continuous channel state 1 (**Fig. 6b-ii top-lateral view, bottom-cross-section; Suppl. Fig. 2h**) to (3) Continuous channel state 2 (**Fig. 6b-iii, top-lateral view, bottom-cross-section; Suppl. Fig. 2j**) and back to Closed discontinuous state (**Fig. 2B-i, 6b-iv, top-lateral view, bottom-cross-section**). The pore volumes of all the putative states of *E-put* respectively (**Fig. 6c**) showed an increase in pore volume during the transition from the closed discontinuous state (**Fig. 6c-i**) to continuous channel states 1 and 2 (**Fig. 6c-ii-iii**). The volume finally decreases when the conformation returns back to the closed discontinuous state (**Fig. 6c-iv).** The water docking phenomena performed in SWISS-DOCK with all the three putative conformational states further validates this hypothesis (**Suppl. Fig. 6a-i-iii**). As a molecular mechanism of how these physical states might be achieved, the orientations of the bottleneck residue PHE 26 in all the three states were analyzed: (1) Closed discontinuous state (**Fig. 6e-i, side view, Suppl. Fig. 6b-i-iii**) (2) Continuous channel state 1 (**Fig. 6e-ii, side view, Suppl. Fig. 6c-i-iii)** (3) Continuous channel state 2 (**Fig. 6e-iii, side view, Suppl. Fig. 6d-i-iii**). We observe a particular recurring change in the PHE 26 orientation during the putative conformational changes from discontinuous closed to the continuous open state and back to the closed discontinuous state (**Fig. 6e-iv, side view, Suppl. Fig. 6b-iii).** On the other hand, the orientation of the PHE 4 residue across the three conformations acts as a gate regulating the putative closed and open states of *E-put* (**Suppl. Fig. 6e**). The conformational changes during its transition from the putative closed to open states in the *E-put* protein could be a biophysical mechanism of regulation of the pore, resembling that of influenza A M2 viroporin (25).

**Figure 6.**
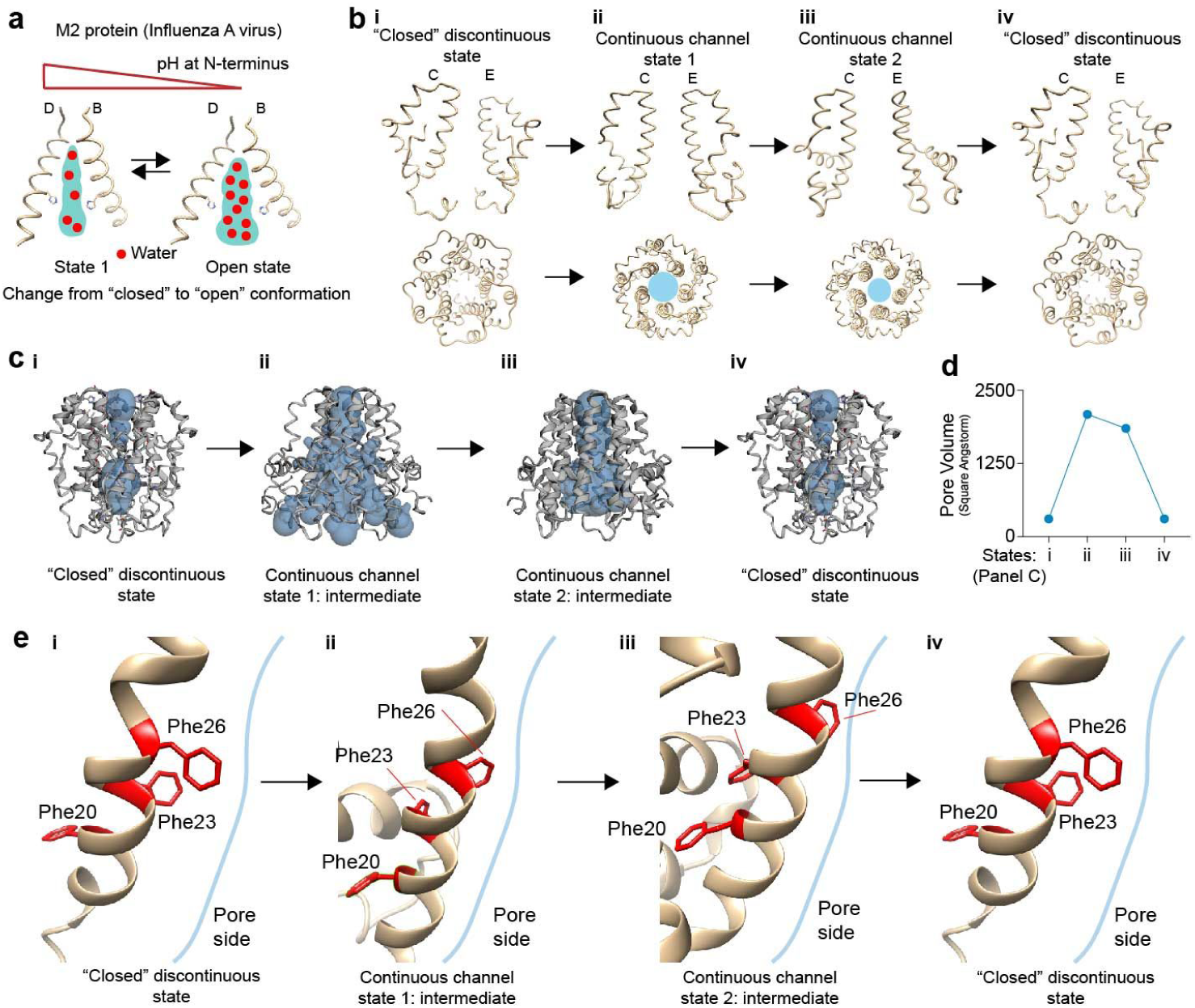
**(a)** Proton channeling by water hopping mechanism in M2 viroporin of influenza A virus in a pH-dependent fashion generated with CHIMERA. **(b)** Different conformational states of *E-put* protein in SARS-CoV-2 by CHIMERA showing its (i) Closed Discontinuous state (ii) Continuous Channel state 1 (iii) Continuous Channel state 2 (iv) Closed Discontinuous state generated with CHIMERA. **(c)** Pore volumes of the *E-put* protein (i) Closed Discontinuous state (ii) Continuous Channel state 1 (iii) Continuous Channel state 2 (iv) Closed Discontinuous state generated with CHIMERA and CASTp. **(d)** Representation of the change of pore volume as in C- i, ii, iii, iv. **(e)** The orientation of the bottleneck PHE 26 in different conformational states as in C- i, ii, iii, iv.

## Discussion

Structural determination of proteins using NMR spectroscopy provided a set of models of the E protein from SARS-CoV-1 (Pdb id: 5×29) having the minimum energies (**Supp. Fig. 6-a-i, side view, Fig. 6-a-ii, top view**). Homology modeling of E-put using this template provided us with many structural models of the protein. Some of them upon proper structural refinement, represent functionally relevant structures despite having a higher Clash score or MolProbity score. The higher scores signify that the particular model is in a higher energy state correlative with functional intermediates of the protein during the viral progression.

We show that the E protein of novel SARS-CoV-2 might assemble on the ER-Golgi network and ERGIC complex of human cells as pentamers similar to previous reports in other coronavirus mediated diseases like SARS-CoV-1 and MERS (19). We have identified various component residues of the pentameric E protein lining its luminal surface which might take part in various molecular events during its conformational changes and corresponding functionalities. The ARG 38 seems to act as the hook of the protein anchoring it to the mammalian ER-Golgi membrane. The orientation of the residues of the hydrophobic funnel (LEU 37, ALA 36, ILE 33) enables water molecules to travel inside the funnel without any critical steric clashes. This is evident from the large distances of their side chains from the docked water molecule, which is beyond the H-bonding limit and short-range interactions limit as well.

The *E-put* protein has a bottleneck located at the midway of the pore and mediated by PHE 26. The bottleneck is followed by a hydrophobic pocket which we call the central core, lined by ALA 22 and LEU 18. ASN 15 and GLU 8 are the polar/charged residues present below the central pore which are at H-bond forming distance from water molecules inside the luminal surface of the pore. The N-terminal end of the *E-put* protein has a gate-like region mediated by PHE 4. The proposed intermediate structures of the *E-put* which were two of the 16 GalaxyWEB outputs, had MolProbity and Clash score in the following order: Closed discontinuous state < Continuous Channel state 1 > Continuous Channel state 2 > Closed discontinuous state. The Continuous Channel state 1 and 2 have less stability and possibly higher energy states than the closed discontinuous state. This change of conformation of *E-put* protein is mediated by the PHE 26 at the bottleneck region which changes its orientation during the transition to the intermediates and PHE 4 which attains closed and open conformations during respective intermediates. The change in pore volume among the conformations also aligns with our hypothesis of intermediate states regulating the *E-put* protein. All these *in silico* findings show that the *E-put* protein may act as a proton channel. The movement of the protons is mediated by the formation of continuous chains of water clusters *via* a water-hopping mechanism partly similar to the M2 proton channel of influenza A virus as reported by Acharya et al., 2010 (25).

In order to gain an understanding of the E protein of novel SARS-CoV-2, we resorted to an *in-silico* approach. It is to be noted that the protein is highly hydrophobic and has a transmembrane domain which increases the difficulty of protein purification and crystallographic studies. However, they can be easily targeted for site-directed mutagenesis or deletion of a functional domain. LAV is a vaccine strategy that takes advantage of the existing virulence of the virus but consists of one or more mutations or deletion of specific regions of the protein resulting in the loss of its pathogenic activity. As a result, the host is able to generate immunological memory to eliminate existing viruses and provide future protection. A possible strategy could be the deletion of the transmembrane domain (12-37) of the *E-put* protein, keeping the ER-Golgi targeting sequence in the C-terminal intact, which might lead to attenuation of the virus. Secondly, site-directed mutagenesis of the residues like ASN 15 and GLU 8 could also eliminate its channeling activity. In addition, mutation of PHE 4 and PHE 26 can render rigidity to the structure affecting its ability to change its conformation during its functionality.

During the recent MERS near-pandemic in and around 2012, a LAV was developed with a MERS coronavirus having truncation near the C-terminal end without disturbing the PDZ-binding motif and some mutations in the NS1 protein of the virus. Keeping that in mind, our study gives a structural insight into the mechanism of action of the novel SARS-CoV-2 E protein as a conformation-dependent proton channel. This presents a potential target for LAV development and other therapeutic strategies using inhibitors.

## Methods

### Multiple sequence alignment

In order to assess the conservation among candidate coronavirus E-proteins, we performed a protein-protein BLAST (BLASTp: Basic Local Alignment Search Tool) with the E protein of SARS-CoV-2 isolated from Shanghai in April (accession number in QII57162.1). Using a 90-100% sequence identity as filters, we aligned the resulting sequences using the online alignment tool Clustal Omega (27) and MegaX (28). Both the software enables high fidelity multiple protein sequence alignment. Jalview (www.jalview.org) was used for visualizing the sequence alignment, conservation score, quality of alignment and consensus residue for each position. The same methods were undertaken for spike proteins (100 sequences), M proteins (54 sequences) and protein 7a (30 sequences).

### Disparity score and mutability

Estimates of evolutionary divergence are shown for the different E-proteins isolated from different sources (bat (HKU3-7), human SARS 2018, bat SARS_RsSHC014, BtRI-BetaCoV, SARS-CoV-1 and SARS-CoV-2 (accession number: QII57162.1)). Analyses were conducted using the Poisson correction model. All ambiguous positions were removed for each sequence pair (pairwise deletion option). The disparity index is a measure of heterogeneity among sites (26) and was calculated from MegaX (28). Briefly, the disparity index uses a Monte-Carlo procedure to test homogeneity among the input sequences. Values greater than 0 indicate the larger differences in amino acid composition biases than expected, based on the evolutionary divergence between sequences and by chance alone.

Frequencies of different amino acids were calculated across the aligned sequences. Mutability was defined as the probability of an amino acid to be non-consistent across the sequence and was calculated as follows:

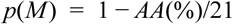

where AA: amino acid composition in percentage and 21 indicates the total amino acid residue space. The same analytical methods for spike proteins were performed. All data is represented as mean ± SD. **Homology modeling of the E-protein:**

The structure of the envelope (E) protein of SARS-CoV-2 was modeled with a template-based homology modeling approach using the NMR structure of E protein from SARS-CoV-1 (PDB id: 5×29;(10)). The full-length pentameric model of the E protein was generated using the following steps:

1. Backbone and side-chain modeling of the monomeric unit.
2. Loop refining of the monomeric unit.
3. Pentamer modeling from the monomer unit.
4. Model refinement and further structural fine-tuning of unreliable structural regions, using the GalaxyWEB server (29).

Another pentameric model of the E protein was generated using the same template (PDB id: 5×29) using the SWISS-MODEL (30, 31). The structures obtained from these modeling servers were validated by scores obtained from the MolProbity (32, 33). The structures were chosen by comparing predominantly the different parameters like percentage of Ramachandran favored and unfavoured residues, percentage of favored and unfavored rotamers, MolProbity score, and Clash Score, validating the quality of the modeled protein.

### Structure visualizations

The structural visualizations of the different models were performed using PYMOL. In addition, we used UCSF-CHIMERA (34) for detailed visualization of residues in the structure and for representation throughout the paper. UCSF-CHIMERA has been referred to as CHIMERA throughout the article. Important amino acid residues lining the inner lumen of the pentameric *E-put* protein were identified. Of particular interest was the PHE 26 where we defined the bottleneck region and this was used to calculate the pore radius for different conformations. The radius of the bottleneck region of the *E-put* protein was calculated with the centroid of the triangle formed by the different orientations of PHE 26 in closed or open conformation. The distance from the vertices of the triangle thus formed from the centroid was then used to calculate the effective radius.

For a scalene triangle having side lengths of a,b and c units, the length of the medians are defined as m_a_, m_b_ and m_c_ units on the respective sides as mentioned in the subscript. The length of the medians are:

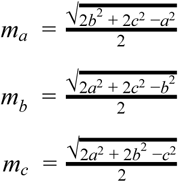

All the medians meet at a point inside the triangle, called the centroid. The distances of the three vertices from the centroid are equal to two-thirds of the length of the median passing through the respective vertices. The distances of each of the vertices from the centroid are considered as the pore radii of the protein in three different directions. The average value of all these radii is calculated as the effective pore radius of the protein.

### Pore volume and topology estimation

The pore volume of the protein in its different conformational states is determined using CASTp (Computed Atlas of Surface Topography of proteins;(35)). In order to account for the presence and impact of the modeled pentameric *E-put* protein, we estimated the presence of surface pockets, internal cavities, and cross channels using the CASTp server. Briefly, CASTp uses a computational geometry approach to measure area and volume and to identify topological imprints on the protein. The lining residues are annotated from the UniProt database and mapped along with the cavity. The cavities with the two maximum molecular surface area and volume were selected. The molecular surface is colored by its hydrophobicity index. These values are determined by the Kyte-Doolittle scale of hydrophobicity with appropriate colors ranging from blue for the most hydrophilic to orange-red for the most hydrophobic pockets. The Ramachandran plots of the protein are plotted using Zeus software (http://www.al-nasir.com/portfolio/; Source: https://www.rcsb.org/pages/thirdparty/molecular_graphics).

### Water molecule interactions in different conformations

The docking of the water molecules to the *E-put* structure was performed in SWISS-DOCK (36).

#### Algorithm

EADock DSS (37) blind docking algorithm is used by SWISS-DOCK and CHARMM (38) force fields are used to create a rank of favorable energies. The most favorable clusters are then used to generate the solvation model.

#### Input files

The modeled protein structure (*E-put*) and ligand (water) were uploaded in the PDB and Mol2 formats respectively. The water molecule (Pubchem ID: CID 962) was imported in CHIMERA in .sdf format and converted to .mol2 format. The SWISS-DOCK server uses CHARMM topology, parameters, and coordinates for determining Merck Molecular Force Field (MMFF) and Van der Waals interactions from the ligand to predict the docking sites on the protein structure.

### Lipid interaction using Autodock VINA

Phosphatidylcholine (Pubchem ID: CID 6441487) and ceramide (Pubchem ID: CID 5702612) are the main lipid components of the ER-Golgi network and ERGIC of mammals including humans. *E-put* has been docked to both the lipid molecules by the following steps:

A. Preparation of lipid molecules in OpenBabel which is an online interface for the preparation of chemical molecules (39).
B. Energy minimization of the modeled structure in CHIMERA using the following steps:
  a. *Fast process:* Steepest descent minimization to remove the critical steric clashes which are highly unfavorable to the protein structure globally.
  b. *Slow process:* Conjugate gradient minimization to remove remaining clashes and other steric discrepancies in the protein structure to reach an energy minimum for that structure.
  c. The energies are reported in kJ/mol in the Reply Log (34).
C. The minimized structure is visualized in CHIMERA and docked to the two lipid molecules using the following steps:
  a. The search space for the docking algorithm to run is defined by a three-dimensional box with appropriate axes’ lengths which encompasses our region of interest in the protein structure.
  b. Docking of the lipid molecules to the E protein in the search space described previously is carried out using Autodock Vina which uses the Broyden-Fletcher-Goldfarb-Shanno (BFGS) algorithm as an Iterated Local search global optimizer method, to speed up its optimization procedures (40).
D. The docked conformations of the protein with the lipid molecules were visualized using CHIMERA and the RMSD values of the docking events analyzed by Vina, are represented by CHIMERA in a Reply Log of Autodock Vina (34, 40).

## Data and materials availability

All data in the main text or the supplementary materials are available upon request.

## Acknowledgments

We thank Mr. Jaydeep Paul for insightful discussions on the research idea, Prof. Félix Rey, Dr. Aleksandra Polosukhina and Dr. Georgios Agoranos for reading the manuscript, suggesting additions and changes, Ms. Madhuparna Chakraborty for proof-reading the manuscript. We earnestly thank Prof. Pierre-Marie Lledo and Prof. Pinak Chakrabarti for providing encouragement to conduct the work and reading the finished manuscript.

## Figures and Tables

**Table 1.** Table showing molprobity score, clash score, and pore radius for three possible conformations of the *E-put* protein.

| Models | Molprobit Score | Clash Score | Pore radius (angstrom) |
| --- | --- | --- | --- |
| E-protein SARS-Cov 2 (Galaxy model) | 0.81 | 0 | 3.4 |
| E-protein SARS-Cov 2 (Galaxy model: open; Suppl Fig 3C) | 2.79 | 15.66 | 5.6 |
| E-protein SARS-Cov 2 (Galaxy model: closed bottleneck; Suppl Fig 3H) | 2.02 | 12.86 | 2.3 |

