## Supplementary Information for "Structural insight into the putative role of novel SARS CoV-2 E protein in viral infection: a potential target for LAV development and therapeutic strategies"

**This PDF file includes:**

Figures S1 to S6

**Other supplementary materials for this manuscript include the following:**

Abbreviations

Supplementary Figure 1

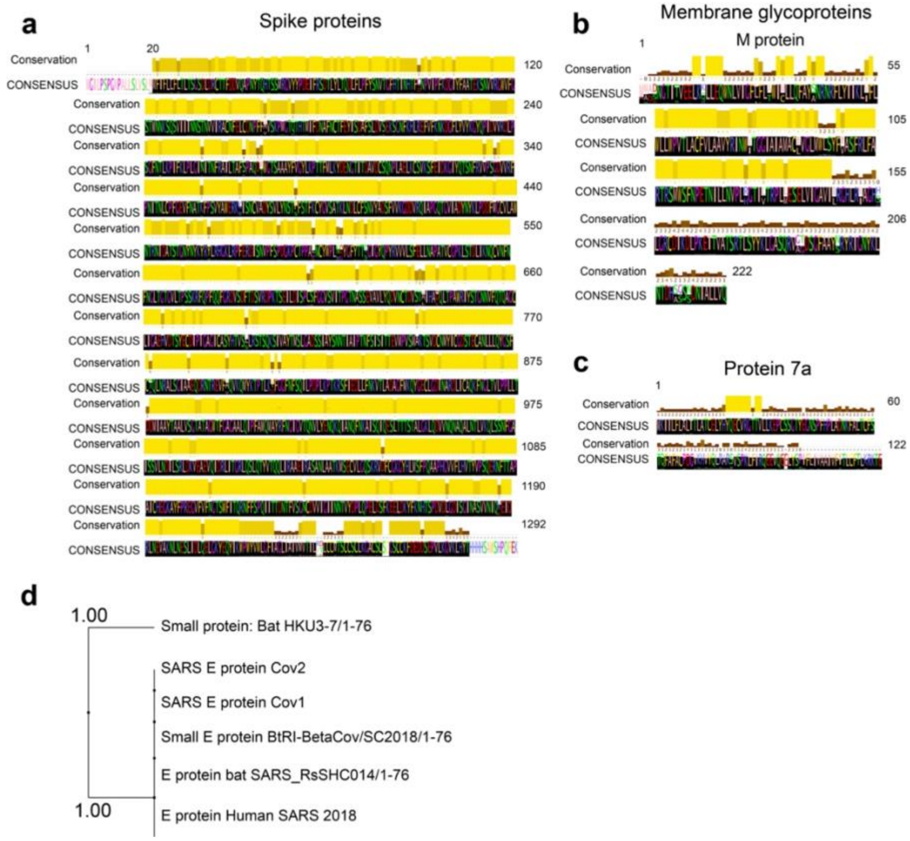


**Supplementary Figure S1: (a)** Conservation score and consensus sequence obtained from multiple sequence alignment for spike protein after a BLASTp search. **(b)** Conservation score and consensus sequence obtained from multiple sequence alignment for M-protein after a BLASTp search. **(c)** Conservation score and consensus sequence obtained from multiple sequence alignment for protein 7a after a BLASTp search. **(d)** Clustering showing close association of the sequences aligned in panel B and their sequence distance.

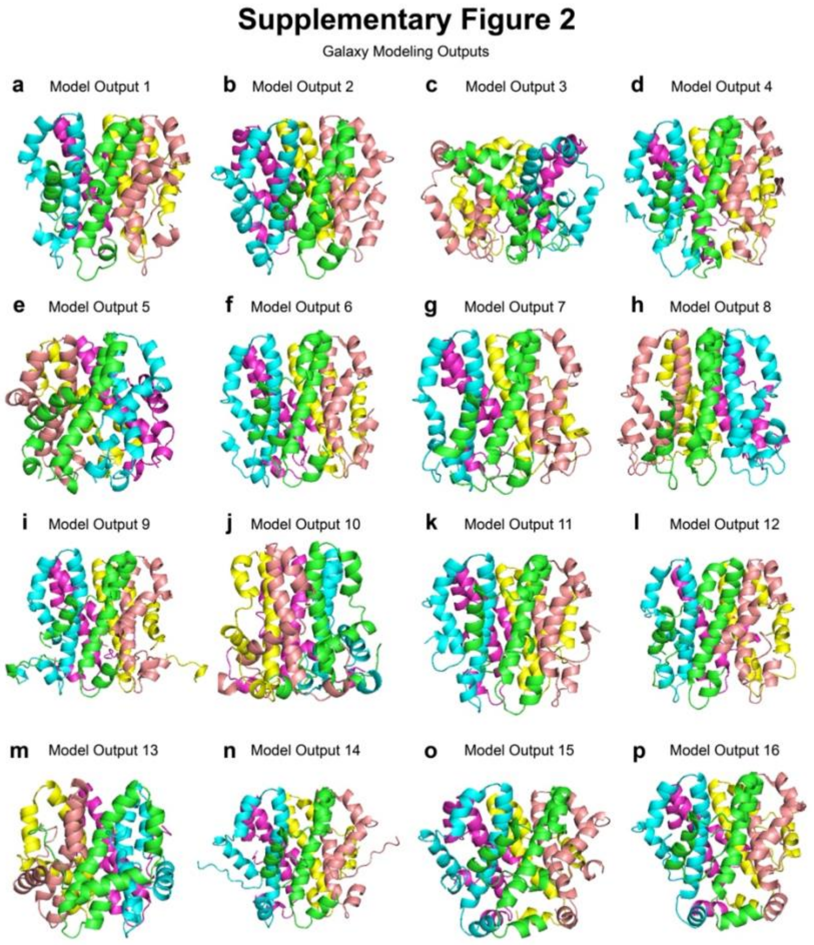


**Supplementary Figure S2: a-p**: 16 output models from GalaxyWEB based on validation parameters from MolProbity analysis.

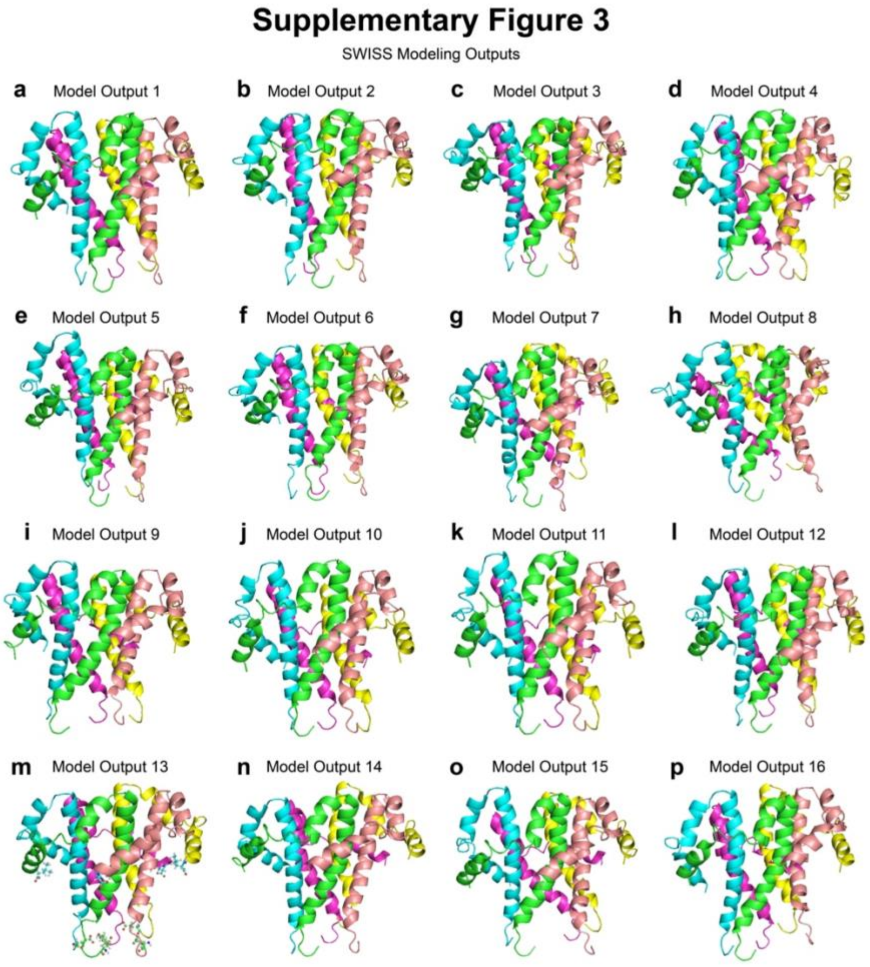


**Supplementary Figure S3: a-p**: 16 output models from SWISS-MODEL based on validation parameters from MolProbity analysis.

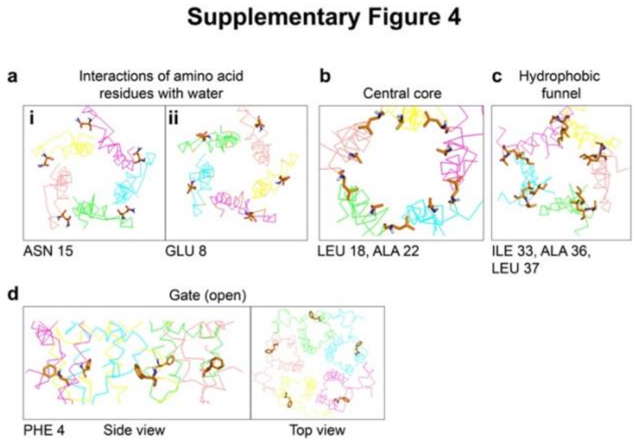


**Supplementary Figure S4: (a)** Top view of residues' orientation of the *E-put* protein generated from PYMOL (i) ASN 15 (ii) GLU 8. **(b)** Top view of residues' orientation of the central core of the *E-put* protein generated from PYMOL. **(c)** Top view of residues' orientation of the hydrophobic funnel of the *E-put* protein generated from PYMOL. **(d)** Residues' orientation generated from PYMOL (i) side view (ii) top view showing the Gate of the *E-put* protein in the open conformation.

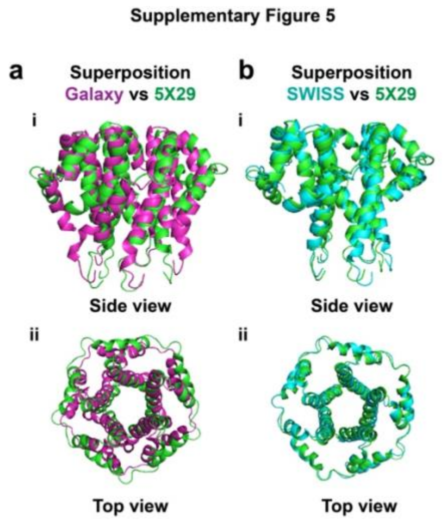


**Supplementary Figure S5: (a)** Superposition of the *E-put* and the 5X29 CoV-1 template protein (purple: *E-put*; green: 5X29) as (i) side view and (ii) top view. **(b)** Superposition of the SWISS-MODEL and the 5X29 CoV-1 template protein (green: 5X29; cyan: SWISS-MODEL) as (i) side view and (ii) top view.

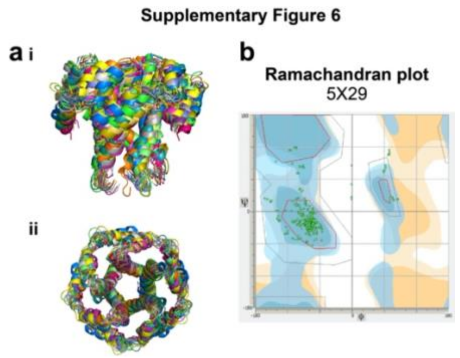


**Supplementary Figure S6: (a)** Structure ensembles of the 5X29 NMR structures as (i) side view and (ii) top view. **(b)** Ramachandran plot of the 5X29 NMR structure of the CoV-1protein (template)

**Supplementary Figure 6**

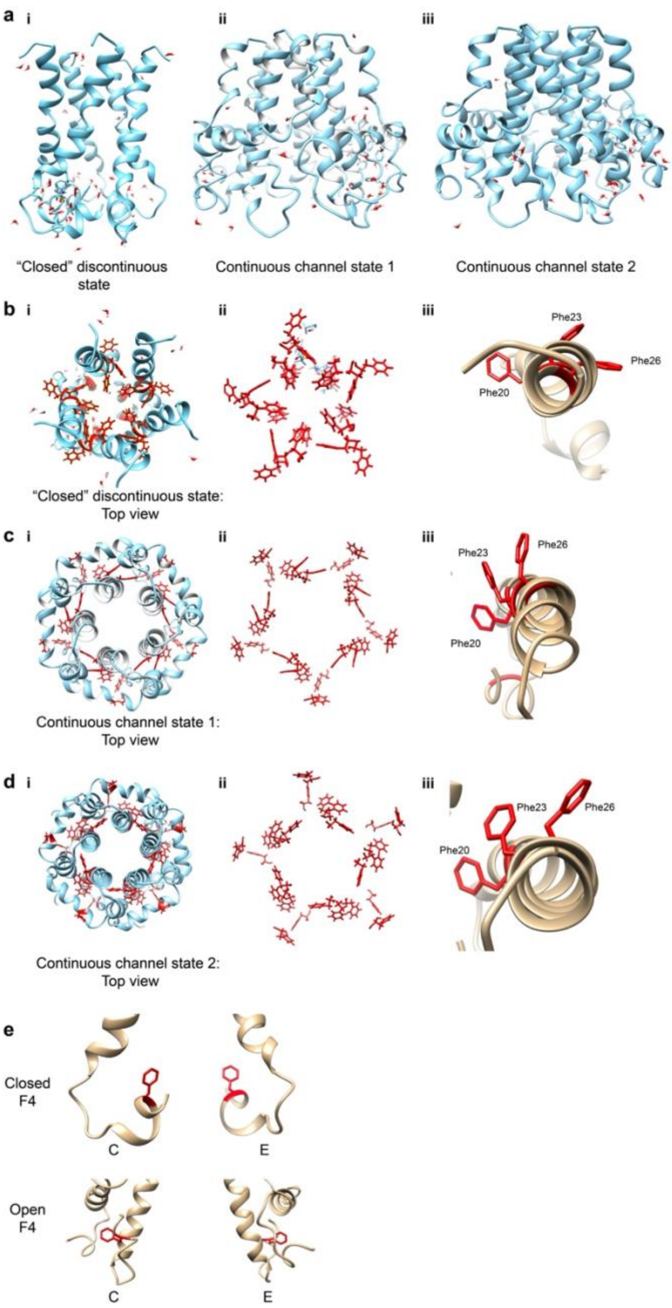


**Supplementary Figure S7: (a)** Docking of water to the modeled structure of *E-put* protein in its different conformations (i) Closed discontinuous state (ii) Continuous channel state 1 (iii) Continuous channel state 2. **(b)** (i-ii) Top view showing all five bottleneck phenylalanine residues (PHE 26) along with adjacent phenylalanine residues (PHE 23 and PHE 20) for the closed discontinuous state. (iii) Enlarged image of the top view orientation of the PHE 26, PHE 23, and PHE 20 on the E chain. **(c)** (i-ii) Top view showing all five bottleneck phenylalanine residues (PHE 26) along with adjacent phenylalanine residues (PHE 23 and PHE 20) for the open continuous channel state 1. (iii) Enlarged image of the top view orientation of the PHE 26, PHE 23, and PHE 20 on the E chain. **(d)** (i-ii) Top view showing all five bottleneck phenylalanine residues (PHE 26) along with adjacent phenylalanine residues (PHE 23 and PHE 20) for the continuous channel state 2. (iii) Enlarged image of the top view orientation of the PHE 26, PHE 23, and PHE 20 on the E chain. **(e)** The orientation of the PHE 4 residues of the Gate of the E protein generated in CHIMERA in its closed and open conformational state.

**Abbreviations:**

Å : Angstrom; ALA : Alanine; ASN : Asparagine; ARG : Arginine; ASP : Aspartic acid; BFGS : Broyden-Fletcher-Goldfarb-Shanno algorithm; BLASTp : Basic local alignment search tool for proteins; CASTp : Computed Atlas of Surface Topography of proteins; COVID-19 : Coronavirus disease 2019; CYS : Cysteine; E protein : Envelope protein; E-put : putative E protein; ER : Endoplasmic Reticulum; ERGIC : ER-Golgi intermediate compartment; GLU : Glutamic Acid; GLY : Glycine; GLN : Glutamine; H-bond : Hydrogen bond; hCoV : human coronavirus; HIS : Histidine; ILE : Isoleucine; LAV : Live attenuated vaccine; LEU : Leucine; LYS : Lysine; MERS : Middle-east respiratory syndrome; MET : Methionine; MMFF : Merck Molecular Force Field; NMR : Nuclear magnetic resonance; NS1 : Non-structural protein 1; PDB : Protein data bank; PPI : Protein-protein interactions; PDZ : Postsynaptic density protein 95 (PSD95)/Drosophila disc large tumor suppressor (Dlg1)/Zonula occludens-1 protein (zo-1); PHE : Phenylalanine; PRO : Proline; RdRp : RNA dependent RNA polymerase; RMSD : Root Mean Square Deviation; RNA : Ribonucleic Acid; SARS : Severe Acute Respiratory Syndrome; SER : Serine; THR : Threonine; TRP : Tryptophan; TYR : Tyrosine; VAL : Valine
